# The SEK-1 p38 MAP kinase pathway modulates Gq signaling in *C. elegans*

**DOI:** 10.1101/093252

**Authors:** Jill M. Hoyt, Samuel K. Wilson, Madhuri Kasa, Jeremy S. Rise, Irini Topalidou, Michael Ailion

## Abstract

Gq is a heterotrimeric G protein that is widely expressed in neurons and regulates neuronal activity. To identify pathways regulating neuronal Gq signaling we performed a forward genetic screen in *Caenorhabditis elegans* for suppressors of activated Gq. One of the suppressors is an allele of *sek-1,* which encodes a mitogen-activated protein kinase kinase (MAPKK) in the p38 MAPK pathway. Here we show that *sek-1* mutants have a slow locomotion rate and that *sek-1* acts in acetylcholine neurons to modulate both locomotion rate and Gq signaling. Furthermore, we find that *sek-1* acts in mature neurons to modulate locomotion. Using genetic and behavioral approaches we demonstrate that other components of the p38 MAPK pathway also play a positive role in modulating locomotion and Gq signaling. Finally, we find that mutants in the SEK-1 p38 MAPK pathway partially suppress an activated mutant of the sodium leak channel NCA-1/NALCN, a downstream target of Gq signaling. Our results suggest that the SEK-1 p38 pathway may modulate the output of Gq signaling through NCA-1.

## Introduction

Gq is a widely expressed heterotrimeric G protein that regulates a variety of biological processes ranging from neurotransmission to cardiovascular pathophysiology (Sánchez-Fernández *et al.* 2014). In the canonical Gq pathway, Gq activates phospholipase Cβ (PLCβ), which cleaves phosphatidylinositol 4,5-bisphosphate (PIP_2_) into the second messengers diacylglycerol (DAG) and inositol trisphosphate (IP_3_) (Rhee 2001). In addition to PLCβ, other Gq effectors have been identified including kinases, such as protein kinase Cζ (PKCζ) and Bruton’s tyrosine kinase (Btk) (Bence *et al.* 1997; García-Hoz *et al.* 2010; Vaqué *et al.* 2013), and guanine nucleotide exchange factors (GEFs) for the small GTPase Rho, such as Trio (Williams *et al.* 2007; Vaqué *et al.* 2013). These noncanonical effectors bridge the activation of Gq to other cellular signaling cascades.

In order to study noncanonical pathways downstream of Gq, we used the nematode *C. elegans* which has a single Gαq homolog (EGL-30) and conservation of the other components of the Gq signaling pathway (Koelle 2016). In neurons, EGL-30 signals through EGL-8 (PLCβ) (Lackner *et al.* 1999) and UNC-73 (ortholog of Trio RhoGEF) (Williams *et al.* 2007). UNC-73 activates RHO-1 (ortholog of RhoA), which has been shown to enhance neurotransmitter release through both diacylglycerol kinase (DGK-1)-dependent and DGK-1-independent pathways (McMullan *et al.* 2006).

To identify additional signaling pathways that modulate Gq signaling, we screened for suppressors of the activated Gq mutant *egl-30(tg26)* (Doi and Iwasaki 2002). *egl-30(tg26)* mutant animals exhibit hyperactive locomotion and a “loopy” posture in which worms have exaggerated, deep body bends and loop onto themselves (Bastiani *et al.* 2003; Topalidou *et al.* 2017). Here we identify one of the suppressors as a deletion allele in the gene *sek-1.* SEK-1 is a mitogen-activated protein kinase kinase (MAPKK), the *C. elegans* ortholog of mammalian MKK3/6 in the p38 MAPK pathway (Tanaka-Hino *et al.* 2002). The p38 MAPK pathway has been best characterized as a pathway activated by a variety of cellular stresses and inflammatory cytokines (Kyriakis and Avruch 2012). However, the p38 MAPK pathway has also been shown to be activated downstream of a G protein-coupled receptor in rat neurons (Huang *et al.* 2004). Btk, a member of the Tec family of tyrosine kinases, has been shown to act downstream of Gq to activate the p38 MAPK pathway (Bence *et al.* 1997), but *C. elegans* lacks Btk and other Tec family members (Plowman *et al.* 1999).

SEK-1 is activated by the MAPKKK NSY-1 (ortholog of ASK1) and activates the p38 MAPKs PMK-1 and PMK-2 (Andrusiak and Jin 2016). The p38 MAPK pathway consisting of NSY-1, SEK-1, and PMK-1 is required for innate immunity in *C. elegans* (Kim *et al.* 2002). NSY-1 and SEK-1 are also required for the specification of the asymmetric AWC olfactory neurons (Sagasti *et al.* 2001; Tanaka-Hino *et al.* 2002); the p38 orthologs PMK-1 and PMK-2 function redundantly in AWC specification (Pagano *et al.* 2015). For both innate immunity and AWC specification, the p38 MAPK pathway acts downstream of the adaptor protein TIR-1 (an ortholog of SARM) (Couillault *et al.* 2004; Chuang and Bargmann 2005). Here we show that the pathway consisting of TIR-1, NSY-1, SEK-1, PMK-1 and PMK-2 also acts to modulate locomotion downstream of Gq signaling.

## Materials and Methods

### C. elegans strains and maintenance

All strains were cultured using standard methods and maintained at 20°C (Brenner 1974). The *sek-1(yak42)* mutant was isolated from an ENU mutagenesis suppressor screen of the activated Gq mutant *egl-30(tg26)* (Ailion *et al.* 2014). *sek-1(yak42)* was outcrossed away from *egl-30(tg26)* before further analysis. Double mutant strains were constructed using standard methods (Fay 2006), often with linked fluorescent markers (Frokjaer-Jensen *et al.* 2014) to balance mutations with subtle visible phenotypes. Table S1 contains all the strains used in this study.

### Mapping

*yak42* was mapped using its slow locomotion phenotype and its *egl-30(tg26)*suppression phenotype. *yak42* was initially mapped to the X chromosome using strains EG1000 and EG1020, which carry visible marker mutations. These experiments showed that *yak42* was linked to *lon-2,* but at least several map units away. *yak42* was further mapped to about one map unit (m.u.) away from the red fluorescent insertion marker *oxTi668* which is located at +0.19 m.u. on the X chromosome.

### Whole-genome sequencing

Strain XZ1233 *egl-30(tg26); yak42* was used for whole-genome sequencing to identify candidate *yak42* mutations. XZ1233 was constructed by crossing a 2X outcrossed *yak42* strain back to *egl-30(tg26).* Thus, in XZ1233, *yak42* has been outcrossed 3X from its original isolate. DNA was isolated from XZ1233 and purified according to the Hobert Lab protocol (http://hobertlab.org/whole-genome-sequencing/). Ion Torrent sequencing was performed at the University of Utah DNA Sequencing Core Facility. The resulting data contained 10,063,209 reads of a mean read length of 144 bases, resulting in about 14X average coverage of the *C. elegans* genome. The sequencing data were uploaded to the Galaxy web platform and we used the public server at usegalaxy.org to analyze the data (Afgan *et al.* 2016). We identified and annotated variants with the Unified Genotyper and SnpEff tools, respectively (DePristo *et al.* 2011; Cingolani *et al.* 2012). We filtered out variants found in other strains we sequenced, leaving us with 605 homozygous mutations. The X chromosome contained 94 mutations: 55 SNPs and 39 indels. Of these, four SNPs were non-synonymous mutations in protein-coding genes, but only two were within 5 m.u. of *oxTi668.* However, we were unable to identify *yak42* from the candidate polymorphisms located near *oxTi668.* Transgenic expression of the most promising candidate *pcyt-1* did not rescue *yak42.* Instead, to identify possible deletions, we scrolled through 2 MB of aligned reads on the UCSC Genome Browser starting at -4.38 m.u. and moving towards the middle of the chromosome (0 m.u.), looking for regions that lacked sequence coverage. We found a 3713 bp deletion that was subsequently confirmed to be the *yak42* causal mutation, affecting the gene *sek-1* located at -1.14 m.u.

### Locomotion assays

Locomotion assay plates were made by seeding 10 cm nematode growth medium plates with 150 μl of an *E. coli* 0P50 stock culture, spread with sterile glass beads to cover the entire plate. Bacterial lawns were grown at room temperature (22.5°C-24.5°C) for 24 hrs and then stored at 4°C until needed. All locomotion assays were performed on first-day adults at room temperature (22.5°C-24.5°C). L4 stage larvae were picked the day before the assay and the experimenter was blind to the genotypes of the strains assayed. For experiments on strains carrying extrachromosomal arrays, the *sek-1(km4)* control worms were animals from the same plate that had lost the array.

Body bend assays were performed as described (Miller *et al.* 1999). A single animal was picked to the assay plate, the plate lid was returned, and the animal allowed to recover for 30 s. Body bends were then counted for one minute, counting each time the worm’s tail reached the minimum or maximum amplitude of the sine wave. All strains in an experiment were assayed on the same assay plate. For experiments with *egl-8, unc-73,* and *rund-1* mutants, worms were allowed a minimal recovery period (until the worms started moving forward, 5 sec maximum) prior to counting body bends.

For the heat shock experiment, plates of first-day adults were parafilmed and heat-shocked in a 34°C water bath for 1 hr. Plates were then un-parafilmed and incubated at 20°C for five hours before performing body bend assays.

Radial locomotion assays were performed by picking animals to the middle of an assay plate. Assay plates were incubated at 20°C for 20 hr and the distances of the worms from the starting point were measured.

Quantitative analysis of the waveform of worm tracks was performed as described (Topalidou *et al.* 2017). Briefly, worm tracks were photographed and Image J was used to measure the period and amplitude. The value for each animal was the average of five period/amplitude ratios.

### C. elegans pictures

Pictures of worms were taken at 60X on a Nikon SMZ18 microscope with the DS-L3 camera control system. The worms were age-matched as first-day adults and each experiment set was photographed on the same locomotion assay plate prepared as described above. The images were processed using ImageJ and were rotated, cropped, and converted to grayscale.

### Molecular biology

Plasmids were constructed using the Gateway cloning system (Invitrogen). Plasmids and primers used are found in Table S2. The *sek-1* cDNA was amplified by RT-PCR from worm RNA and cloned into a Gateway entry vector. To ensure proper expression of *sek-1,* an operon GFP was included in expression constructs with the following template: (promoter)p::*sek-1*(cDNA)*::tbb-2utr::gpd-2 operon::GFP::H2B:cye-1utr* (Frøkjær-Jensen *et al.* 2012). This resulted in untagged SEK-1, but expression could be monitored by GFP expression.

### Injections

C. elegans strains with extrachromosomal arrays were generated by standard methods (Mello *et al.* 1991). Injection mixes were made with a final total concentration of 100 ng/μL DNA. Constructs were injected at 5 ng/μL, injection markers at 5 ng/μL, and the carrier DNA Litmus 38i at 90 ng/μL. Multiple lines of animals carrying extrachromosomal arrays were isolated and had similar behaviors as observed by eye. The line with the highest transmittance of the array was assayed.

### Statistical analysis

At the beginning of the project, a power study was conducted on pilot body bend assays using wild type and *sek-1(yak42)* worms. To achieve a power of 0.95, it was calculated that 17 animals should be assayed per experiment. Data were analyzed to check if normally distributed (using the D’Agostino-Pearson and Shapiro-Wilk normality tests) and then subjected to the appropriate analysis using GraphPad Prism 5. For data sets with three or more groups, if the data were normal they were analyzed with a one-way ANOVA; if not, with a Kruskal-Wallis test. Post-hoc tests were used to compare data sets within an experiment. Reported p-values are corrected. Table S3 contains the statistical tests for each experiment. p<0.05 = *; p<0.01 = **; p<0.001 = ***.

### Reagent and Data Availability

Strains and plasmids are shown in Table S1 and Table S2 and are available from the *Caenorhabditis* Genetics Center (CGC) or upon request. The authors state that all data necessary for confirming the conclusions presented in the article are represented fully within the article and Supplemental Material.

## Results

### *sek-1* suppresses activated Gq

To identify genes acting downstream of Gαq, we performed a forward genetic screen for suppressors of the activated Gq mutant, *egl-30(tg26)* (Doi and Iwasaki 2002). *egl-30(tg26)* worms are hyperactive and have a “loopy” posture characterized by an exaggerated waveform (Figure 1B-1E). Thus, we screened for worms that are less hyperactive and less loopy. We isolated a recessive suppressor, *yak42,* and mapped it to the middle of the X chromosome (see Materials and Methods). Whole-genome sequencing revealed that *yak42* carries a large deletion of the *sek-1* gene from upstream of the start codon into exon 4 (Figure 1A). *yak42* also failed to complement *sek-1(km4),* a previously published *sek-1* deletion allele, for the Gq suppression phenotype (Figure 1A) (Tanaka-Hino *et al.* 2002).

**Figure 1.**
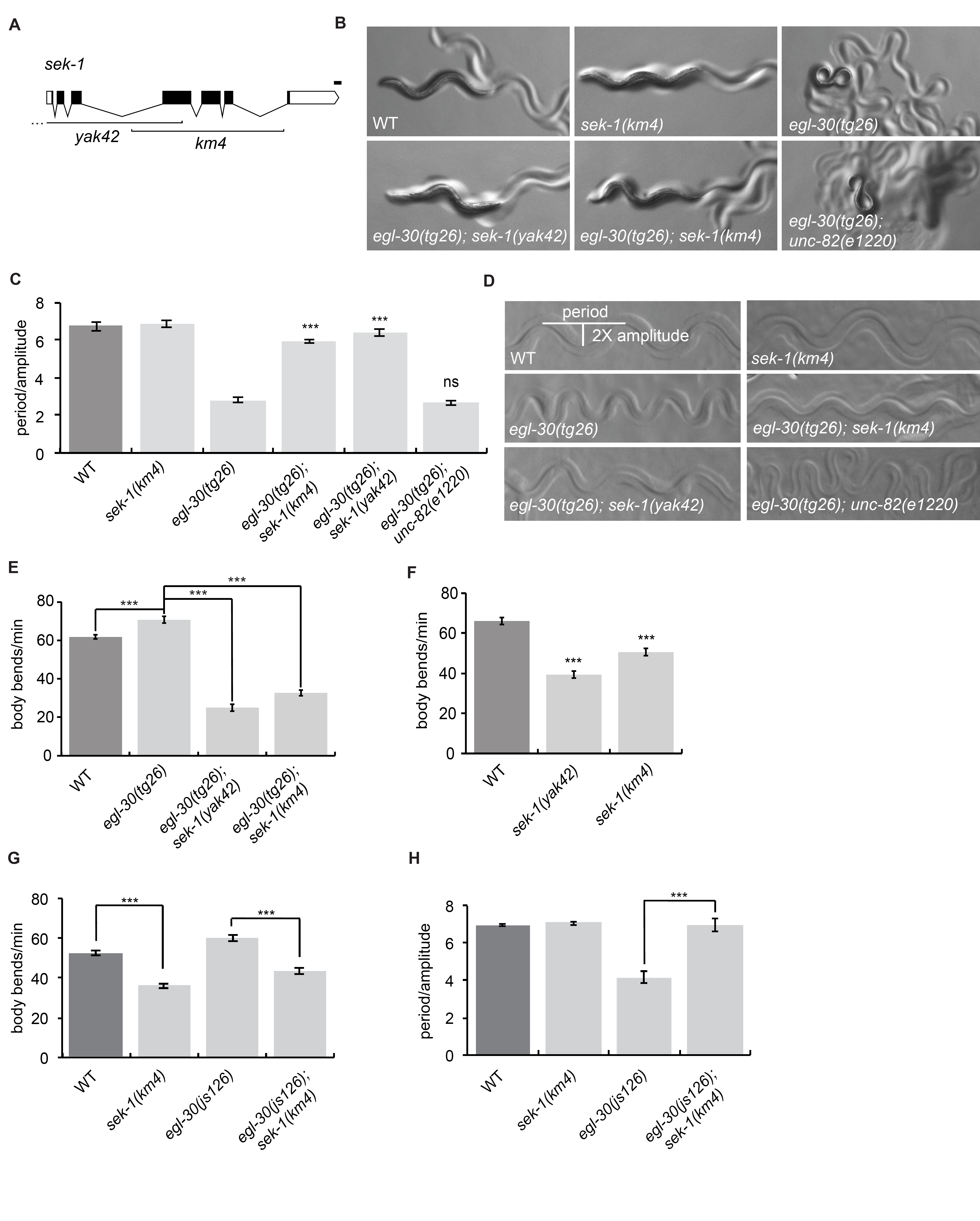
*sek-1* acts downstream of Gαq to modulate locomotion behavior. (A) Gene structure of *sek-1.* White boxes depict the 5’ and 3’ untranslated regions, black boxes depict exons, and lines show introns. The positions of the *yak42* and *km4* deletions are shown. *yak42* is a 3713 bp deletion that extends to 1926 bp upstream of the start codon. Drawn with Exon-Intron Graphic Maker (http://www.wormweb.org/exonintron). Scale bar is 100 bp. (B-D) *sek-1(yak42)* and *sek-1(km4)* suppress the loopy waveform of the activated Gq mutant *egl-30(tg26). unc-82(e1220)* does not suppress *egl-30(tg26).* (B) Photos of first-day adult worms. WT: wild type. (C) Quantification of the waveform phenotype. ***, p<0.001; ns, p>0.05 compared to *egl-30(tg26).* Error bars = SEM, n=5. (D)Photos of worm tracks. (E) The activated Gq mutant *egl-30(tg26)* is hyperactive and is suppressed by *sek-1(yak42)* and *sek-1(km4).* ***, p<0.001, error bars = SEM, n=20. (F) *sek-1* mutant worms have slow locomotion. ***, p<0.001 compared to wild-type. Error bars = SEM, n=20. (G) *sek-1(km4)* suppresses the hyperactive locomotion of the activated Gq mutant *egl-30(js126).* ***, p<0.001, error bars = SEM, n=20. (H) *sek-1(km4)* suppresses the loopy waveform of the activated Gq mutant *egl-30(js126).* ***, p<0.001, error bars = SEM, n=5.

*egl-30(tg26)* double mutants with either *sek-1(yak42)* or *sek-1(km4)* are not loopy (Figure 1B-1D) and are not hyperactive (Figure 1E and S1A). *sek-1(yak42)* was outcrossed from *egl-30(tg26)* and assayed for locomotion defects. Both the *sek-1(yak42)* and *sek-1(km4)* mutants are coordinated but move more slowly than wild-type (Figure 1F). The *sek-1(ag1)* point mutation (Kim *et al.* 2002) also causes a similar slow locomotion phenotype (Figure S1B). To test whether the *egl-30(tg26)* suppression phenotype might be an indirect effect of the slow locomotion of a *sek-1* mutant, we built an *egl-30(tg26)* double mutant with a mutation in *unc-82,* a gene required for normal muscle structure. *unc-82* mutants are coordinated but move slowly, similar to a *sek-1* mutant (Hoppe *et al.* 2010). However, although an *egl-30(tg26) unc-82(e1220)* double mutant moves more slowly than *egl-30(tg26)* (Figure S1C), it is still loopy (Figure 1B-1D). Thus, *sek-1* appears to be a specific suppressor of activated *egl-30.*

The *egl-30(tg26)* allele causes an R243Q missense mutation in the Gα switch III region that has been shown to reduce both the intrinsic GTPase activity of the G protein and render it insensitive to GTPase-activation by a regulator of G protein signaling (RGS) protein, thus leading to increased G protein activation (Natochin and Artemyev 2003). To test whether the suppression of *egl-30(tg26)* by *sek-1* is specific for this *egl-30* allele, we built a double mutant between *sek-1(km4)* and the weaker activating mutation *egl30(js126). egl-30(js126)* causes a V180M missense mutation in the Gα switch I region immediately adjacent to one of the key residues required for GTPase catalysis (Hawasli *et al.* 2004). Thus, the *tg26* and *js126* alleles activate EGL-30 through different mechanisms. The *sek-1(km4)* mutant also suppresses the hyperactivity and loopy waveform of *egl-30(js126)* (Figure 1G, 1H), demonstrating that *sek-1* suppression of activated *egl-30* is not allele-specific.

EGL-30/Gαq is negatively regulated by GOA-1, the worm Gαo/i ortholog, and the RGS protein EAT-16 (Hajdu-Cronin *et al.* 1999). We tested whether *sek-1* also suppresses the *goa-1* and *eat-16* loss-of-function mutants that cause a hyperactive and loopy phenotype similar to activated *egl-30* mutants. *sek-1(km4)* suppresses the hyperactivity and loopy waveform of *goa-1(sa734)* (Figure S1D, S1E). However, though *sek-1(km4)* suppresses the hyperactivity of *eat-16(tm775),* it did not significantly suppress the loopy waveform (Figure S1F, S1G). One possible downstream effector of GOA-1 is the DAG kinase DGK-1 that inhibits DAG-dependent functions such as synaptic vesicle release (Nurrish *et al.* 1999; Miller *et al.* 1999). *dgk-1(sy428)* animals are hyperactive, but the *sek-1 dgk-1* double mutant is uncoordinated and looks like neither *sek-1* nor *dgk-1* mutants, confounding the interpretation of how *sek-1* genetically interacts with *dgk-1.*

### *sek-1* acts in mature acetylcholine neurons

*egl-30* is widely expressed and acts in neurons to modulate locomotion (Lackner *et al.* 1999), so it is possible that *sek-1* also acts in neurons to modulate Gq signaling. *sek-1* is expressed in neurons, intestine, and several other tissues (Tanaka-Hino *et al.* 2002) and has been shown to function in GABA neurons to promote synaptic transmission (Vashlishan *et al.* 2008).

To identify the cell type responsible for the *sek-1* locomotion phenotypes, we expressed the wild-type *sek-1* cDNA under different cell-specific promoters and tested for transgenic rescue of a *sek-1* null mutant. Expression of *sek-1* in all neurons (using the *unc-119* promoter) or in acetylcholine neurons *(unc-17* promoter) was sufficient to rescue the *sek-1* mutant slow locomotion phenotype, but expression in GABA neurons *(unc-47* promoter) was not sufficient to rescue (Figure 2A, B). These results indicate that *sek-1* acts in acetylcholine neurons to modulate locomotion rate.

**Figure 2.**
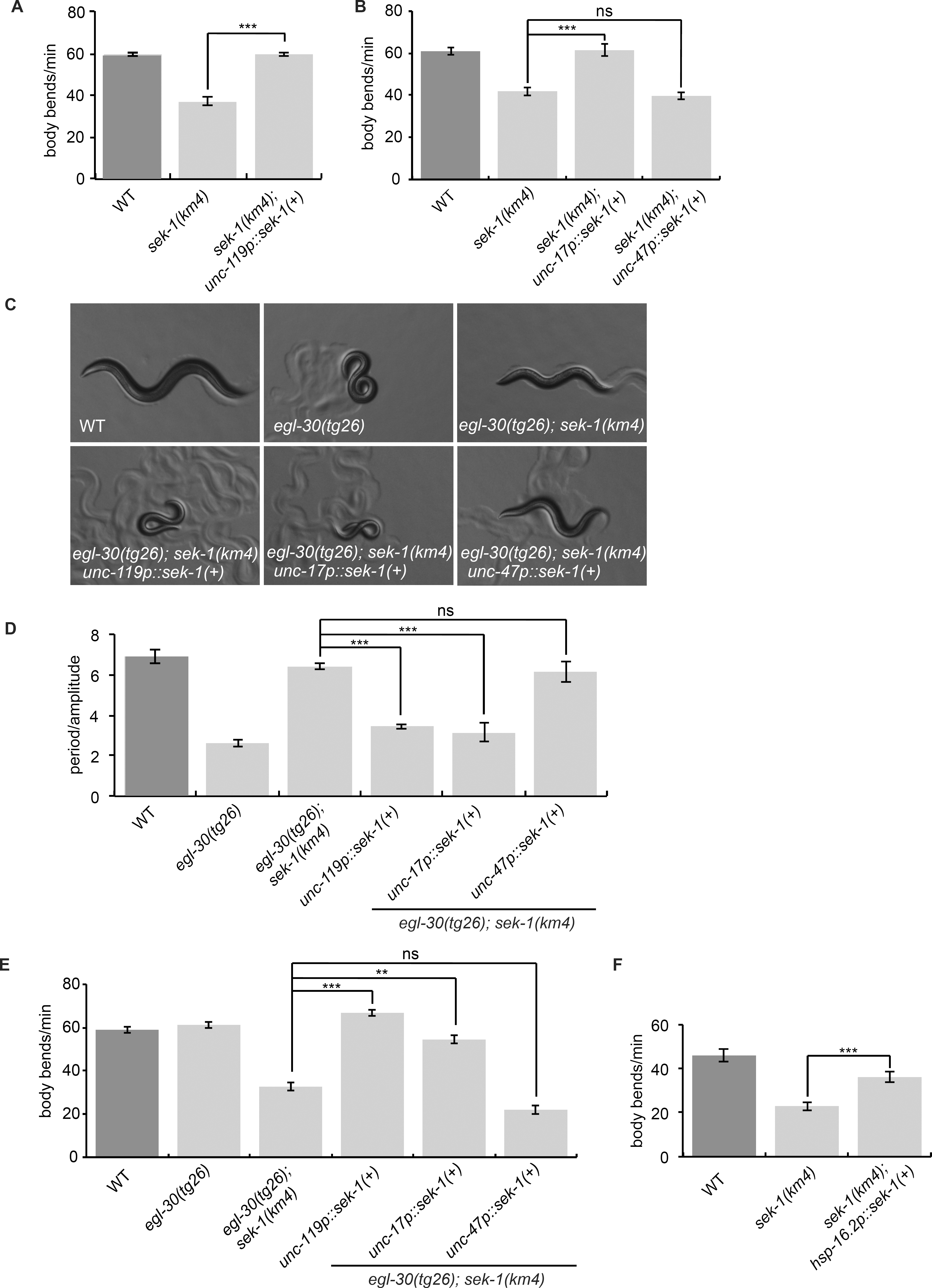
*sek-1* acts in mature acetylcholine neurons to modulate locomotion. (A) *sek-1* acts in neurons to modulate locomotion rate. The *sek-1* wild-type cDNA driven by the *unc-119* pan-neuronal promoter *[unc-119p::sek-1(+)]* rescues the slow locomotion phenotype of *sek-1(km4)* worms. ***, p< 0.001, error bars = SEM, n=20. (B) *sek-1* acts in acetylcholine neurons to modulate locomotion rate. *sek-1* WT cDNA driven by the *unc-17* acetylcholine neuron promoter *[unc-17p::sek-1(+)]* rescues the slow locomotion phenotype of *sek-1(km4)* worms but *sek-1* expression in GABA neurons using the *unc-47* promoter [*unc-47p::sek-1(+)*] does not. ***, p< 0.001, error bars = SEM, n=20. (C-D) *sek-1* acts in acetylcholine neurons to modulate the loopy waveform of *egl-30(tg26). egl-30(tg26) sek-1(km4)* worms expressing *unc-119p::sek-1(+)* or *unc-17p::sek-1(+)* are loopy like *egl-30(tg26),* but *egl-30(tg26) sek-1(km4)* worms expressing *unc-47p::sek-1(+)* are similar to *egl-30(tg26) sek-1.* (C) Photos of worms. (D) Quantification of waveform phenotype. ***, p<0.001; ns, p>0.05. Error bars = SEM, n=5. (E) *sek-1* acts in acetylcholine neurons to modulate the locomotion rate of *egl-30(tg26). egl-30(tg26) sek-1(km4)* worms expressing *unc-119p::sek-1(+)* or *unc-17p::sek-1(+)* have an increased locomotion rate compared to *egl-30(tg26) sek-1,* but *egl-30(tg26) sek-1(km4)* worms expressing *unc-47p::sek-1(+)* are similar to *egl-30(tg26) sek-1.* ***, p< 0.001, **, p<0.01; ns, p>0.05. Error bars = SEM, n=17-20. (F) *sek-1* acts in mature neurons to modulate locomotion rate. Heat-shock induced expression of *sek-1* in adults *(hsp-16.2p::sek-1(+))* rescues the slow locomotion phenotype of *sek-1(km4).* ***, p< 0.001, error bars = SEM, n=20.

We next tested whether *sek-1* acts in neurons to suppress *egl-30(tg26).* Expression of *sek-1* under pan-neuronal and acetylcholine neuron promoters reversed the *sek-1* suppression of *egl-30(tg26).* Specifically, *egl-30(tg26) sek-1* double mutants expressing wild-type *sek-1* in all neurons or acetylcholine neurons resembled the *egl-30(tg26)* single mutant (Figure 2C-E). However, expression of *sek-1* in GABA neurons did not reverse the suppression phenotype (Figure 2C-E). Together, these data show that *sek-1* acts in acetylcholine and not GABA neurons to modulate both wild-type locomotion rate and to modulate Gq signaling.

To narrow down the site of *sek-1* action, we expressed *sek-1* in head *(unc-17H* promoter) and motorneuron *(unc-17β* promoter) acetylcholine neuron subclasses (Topalidou *et al.* 2017). Expression of *sek-1* in acetylcholine motorneurons rescued the *sek-1* slow locomotion phenotype (Figure S2A), suggesting that the slow locomotion of *sek-1* mutants is due to a loss of *sek-1* in acetylcholine motorneurons. However, expression of *sek-1* in either the head acetylcholine neurons or motorneurons partially reversed the *sek-1* suppression of *egl-30(tg26)* hyperactivity (Figure S2B), suggesting that the hyperactivity of activated Gq mutants may result from excessive Gq signaling in both head acetylcholine neurons and acetylcholine motorneurons; *sek-1* may act in Gq signaling in both neuronal cell types. By contrast, expression of *sek-1* in head acetylcholine neurons but not motorneurons reversed the *sek-1* suppression of the *egl-30(tg26)* loopy waveform (Figure S2C), suggesting that the loopy posture of activated Gq mutants may result from excessive Gq signaling in head acetylcholine neurons, and *sek-1* may act in those neurons to control body posture.

Because *sek-1* acts in the development of the AWC asymmetric neurons, we asked whether *sek-1* also has a developmental role in modulating locomotion by testing whether adult-specific *sek-1* expression (driven by a heat-shock promoter) is sufficient to rescue the *sek-1* mutant. We found that *sek-1* expression in adults rescues the *sek-1* slow locomotion phenotype (Figure 2F). This result indicates that *sek-1* is not required for development of the locomotion circuit and instead acts in mature neurons to modulate locomotion.

### The p38 MAPK pathway is a positive regulator of Gq signaling

SEK-1 is the MAPKK in the p38 MAPK pathway consisting of the adaptor protein TIR-1, NSY-1 (MAPKKK), SEK-1 (MAPKK), and PMK-1 or PMK-2 (MAPKs) (Tanaka-Hino *et al.* 2002; Andrusiak and Jin 2016). We tested whether the entire p38 MAPK signaling module also modulates locomotion rate and suppression of activated Gq. Both *tir-1(tm3036)* and *nsy-1(ok593)* mutant animals have slow locomotion on their own and also suppress the hyperactivity and loopy waveform of *egl-30(tg26)* (Figure 3A-D, G and H). We also tested single mutants in each of the three worm p38 MAPK genes *(pmk-1, pmk-2* and *pmk-3)* and a *pmk-2 pmk-1* double mutant. Although we found that the *pmk-2* and *pmk-3* single mutants were slightly slow on their own, only the *pmk-2 pmk-1* double mutant phenocopied *sek-1* and suppressed both the hyperactivity and loopy waveform of *egl-30(tg26)* (Figure 3E-H). Thus, *pmk-2* and *pmk-1* act redundantly downstream of *sek-1* to suppress *egl-30(tg26).* These data suggest that the p38 MAPK pathway modulates locomotion rate in *C. elegans* and acts genetically downstream of *egl-30*.

**Figure 3.**
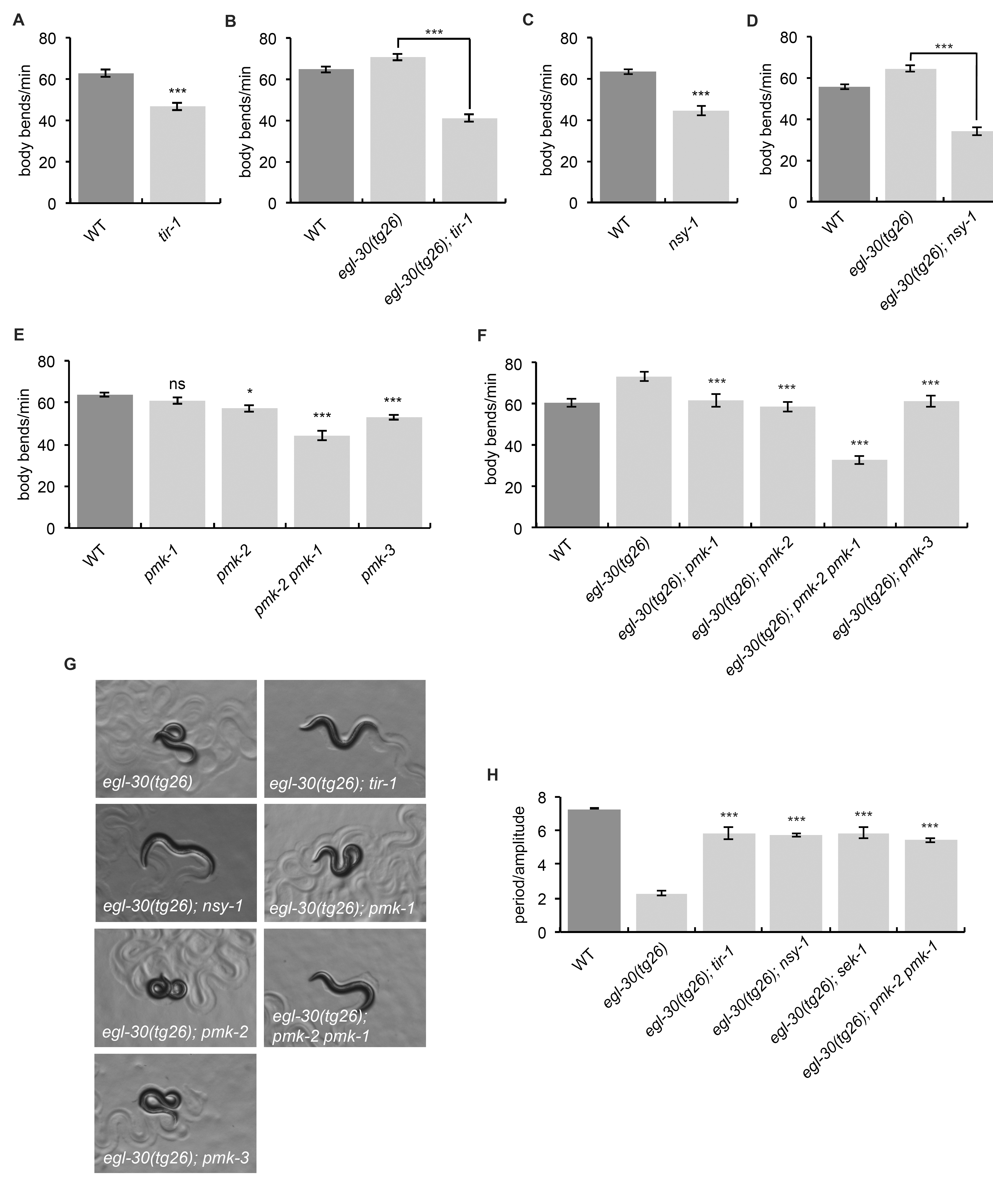
The p38 MAPK pathway modulates locomotion downstream of *egl-30*. (A) *tir-1(tm3036)* mutant animals have slow locomotion. ***, p<0.001, error bars = SEM, n=20. (B) *tir-1(tm3036)* suppresses *egl-30(tg26). egl-30(tg26) tir-1* animals move more slowly than the hyperactive *egl-30(tg26)* animals. ***, p< 0.001, error bars = SEM, n=20. (C) *nsy-1(ok593)* mutant animals have slow locomotion. ***, p<0.001, error bars = SEM, n=20. (D) *nsy-1(ok593)* suppresses *egl-30(tg26). egl-30(tg26) nsy-1* animals move more slowly than the hyperactive *egl-30(tg26)* animals. ***, p< 0.001, error bars = SEM, n=20. (E) *pmk-2, pmk-2 pmk-1,* and *pmk-3* mutant animals have slow locomotion. ***, p< 0.001; *, p<0.05, compared to WT. Error bars = SEM, n=20. (F) A *pmk-2 pmk-1* double mutant suppresses the hyperactivity of *egl-30(tg26).* ***, p< 0.001 compared to *egl-30(tg26).* Error bars = SEM, n=20. (G, H) *tir-1(tm3036), nsy-1(ok593),* and the *pmk-2 pmk-1* double mutant suppress the loopy waveform of *egl-30(tg26). egl-30(tg26)* animals with mutations in either *pmk-1, pmk-2,* or *pmk-3* are still loopy. (G) Worm photos. (H) Quantification. ***, p<0.001 compared to *egl-30(tg26).* Error bars = SEM, n=5.

The JNK MAPK pathway, related to the p38 MAPK family, also modulates locomotion in *C. elegans.* Specifically, the JNK pathway members *jkk-1* (JNK MAPKK) and *jnk-1* (JNK MAPK) have been shown to act in GABA neurons to modulate locomotion (Kawasaki 1999). We found that the *jkk-1* and *jnk-1* single mutants had slow locomotion and that the double mutants with p38 MAPK pathway members exhibited an additive slow locomotion phenotype (Figure S3A). Moreover, neither *jkk-1* nor *jnk-1* suppressed the loopy phenotype of *egl-30(tg26)* (Figure S3B). Thus, the JNK and p38 MAPK pathways modulate locomotion independently and the JNK pathway is not involved in Gq signaling.

We also tested the involvement of possible p38 MAPK pathway effectors. One of the targets of PMK-1 is the transcription factor ATF-7 (Shivers *et al.* 2010). Both the *atf-7(qd22 qd130)* loss-of-function mutant and the *atf-7(qd22)* gain-of-function mutant moved slowly compared to wild-type animals (Figure S3C). However, *atf-7(qd22 qd130)* did not suppress the loopiness of *egl-30(tg26)* (Figure S3B), suggesting that *atf-7* is not a target of this pathway, or else it acts redundantly with other downstream p38 MAPK targets. We also tested *gap-2,* the closest *C. elegans* homolog of ASK1-interacting Protein (AIP1) which activates ASK1 (the ortholog of *C. elegans* NSY-1) in mammalian systems (Zhang *et al.* 2003). A *C. elegans gap-2* mutant has no locomotion defect (Figure S3D). Finally, we tested VHP-1, a phosphatase for p38 and JNK MAPKs that inhibits p38 MAPK signaling (Kim *et al.* 2004). However, the *vhp-1(sa366)* mutant also has no locomotion defect (Figure S3D).

*egl-30(tg26)* animals are loopy and hyperactive so we tested whether increased activation of the TIR-1/p38 MAPK signaling module causes similar phenotypes. The *tir-1(ky648tg26)* allele leads to a gain-of-function phenotype in the AWC neuron specification (Chang *et al.* 2011), but does not cause loopy or hyperactive locomotion (Figure S3E, F).

### Genetic interactions of *sek-1* with pathways acting downstream of Gq

Our forward genetic screen for suppressors of *egl-30(tg26)* identified mutants that fall into three different categories: mutants in the canonical Gq pathway such as the PLC *egl-8* ((Lackner *et al.* 1999), mutants in the RhoGEF Trio pathway such as *unc-73* (Williams et al., 2007), and mutants that affect dense-core vesicle biogenesis and release (Ailion *et al.* 2014; Topalidou *et al.* 2016).

To test if *sek-1* acts in any of these pathways we built double mutants between *sek-1* and members of each pathway. Loss-of-function alleles of *egl-8(sa47), unc-73(ox317),* and *rund-1(tm3622)* have slow locomotion (Figure 4A-C). We found that *sek-1* enhances the slow locomotion phenotype of *egl-8* and *rund-1* single mutants, suggesting that *sek-1* does not act in the same pathway as *egl-8* or *rund-1* (Figure 4A, B). By contrast, *sek-1* does not enhance the slow locomotion phenotype of *unc-73* mutants (Figure 4C), suggesting that *sek-1* may act in the same genetic pathway as the Trio RhoGEF *unc-73.*

**Figure 4.**
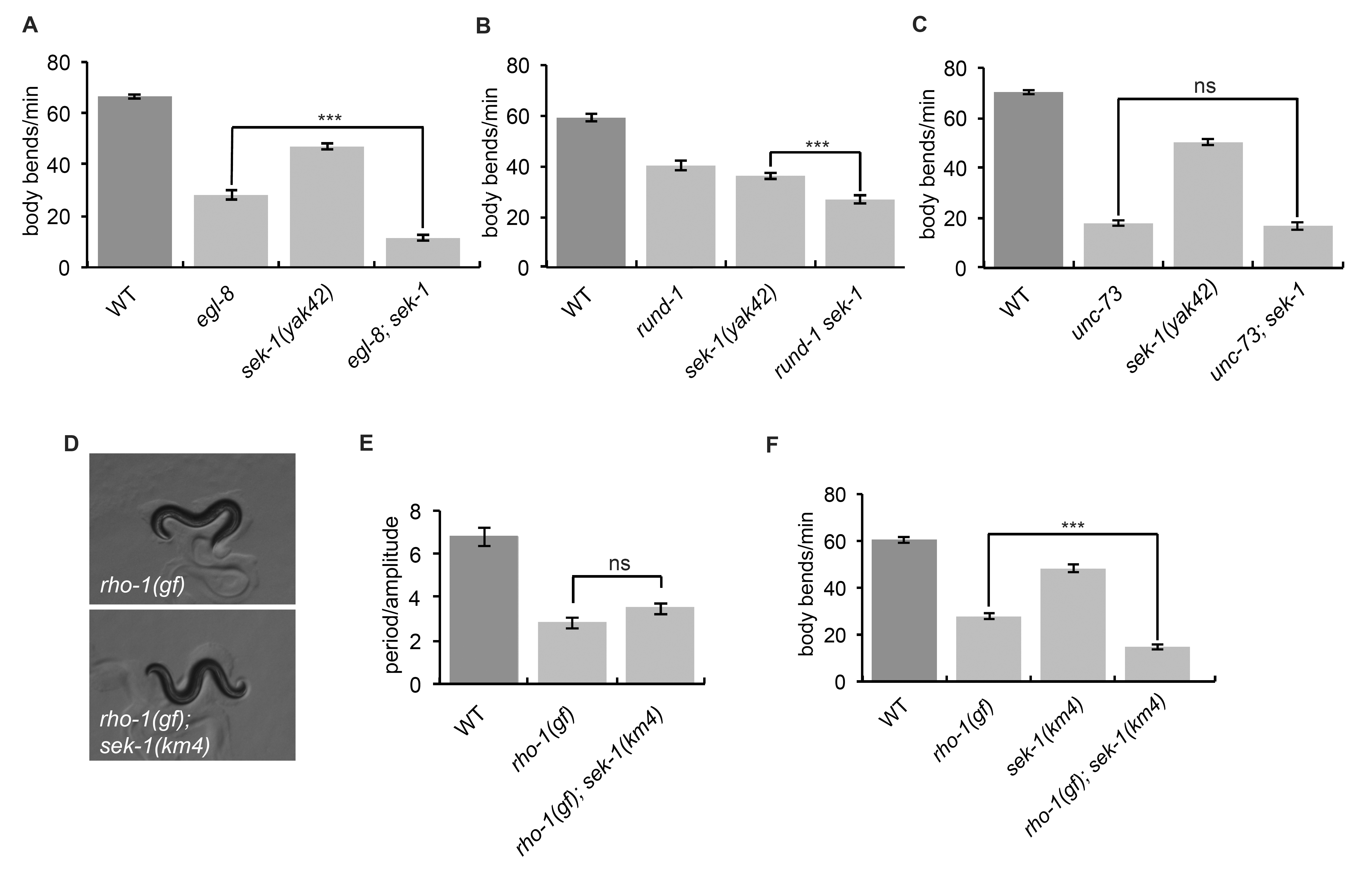
*sek-1* acts in the same genetic pathway as *unc-73*. (A) *sek-1* does not act in the same genetic pathway as *egl-8.* The *sek-1(yak42)* mutation enhances the slow locomotion of the *egl-8(sa47)* mutant. ***, p< 0.001, error bars = SEM, n=20. (B) *sek-1* does not act in the same genetic pathway as *rund-1.* The *sek-1(yak42)* mutation enhances the slow locomotion of the *rund-1(tm3622)* mutant. ***, p< 0.001, error bars = SEM, n=20. (C) *sek-1* may act in the same genetic pathway as *unc-73.* The *sek-1(yak42)* mutation does not enhance the slow locomotion phenotype of the *unc-73(ox317)* mutant. ns, p>0.05, error bars = SEM, n=20. (D-E) *sek-1(km4)* does not suppress the loopy waveform of *nzIs29 rho-1(gf)* animals. (D) Worm photos. (E) Quantification. ns, p>0.05, error bars = SEM, n=5. (F) *sek-1(km4)* does not suppress the slow locomotion of *rho-1(gf)* animals. ***, p< 0.001, error bars = SEM, n=20.

We next tested whether *sek-1* interacts with *rho-1,* encoding the small G protein Rho that is activated by Trio. Because *rho-1* is required for viability (Jantsch-Plunger *et al.* 2000), we used an integrated transgene overexpressing an activated *rho-1* mutant allele specifically in acetylcholine neurons. Animals carrying this activated RHO-1 transgene, referred to here as *rho-1(gf),* have a loopy posture reminiscent of *egl-30(tg26)* (McMullan *et al.* 2006), and a decreased locomotion rate (Figure 4D-F). *rho-1(gf) sek-1(km4)* double mutants had a loopy body posture like *rho-1(gf)* and an even slower locomotion rate (Figure 4D-F), suggesting that *sek-1* and *rho-1(gf)* mutants have additive locomotion phenotypes. However, both *sek-1(km4)* and *sek-1(yak42)* weakly suppress the slow growth rate of the *rho-1(gf)* mutant (data not shown). Because *sek-1* does not enhance *unc-73* mutants and suppresses some aspects of the *rho-1(gf)* mutant, *sek-1* may modulate output of the Rho pathway, though it probably is not a direct transducer of Rho signaling.

### *sek-1* and *nsy-1* partially suppress activated NCA

To clarify the relationship of the SEK-1 p38 MAPK pathway to the Rho pathway acting downstream of Gq, we examined interactions with *nca-1,* a downstream target of the Gq-Rho pathway (Topalidou *et al.* 2017). NCA-1 and its orthologs are sodium leak channels associated with rhythmic behaviors in several organisms (Nash *et al.* 2002; Lu *et al.* 2007; Shi *et al.* 2016). In *C. elegans,* NCA-1 potentiates persistent motor circuit activity and sustains locomotion (Gao *et al.* 2015).

We tested whether *sek-1* and *nsy-1* mutants suppress the activated NCA-1 mutant *ox352,* referred to as *nca-1(gf).* The *nca-1(gf)* animals are coiled and uncoordinated; thus, it is difficult to measure their locomotion rate by the body bend assay because they do not reliably propagate sinusoidal waves down the entire length of their body. Instead, we used a radial locomotion assay in which we measured the distance animals moved from the center of a plate. *nca-1(gf)* double mutants with either *sek-1(km4)* or *nsy-1(ok593)* uncoil a bit but still exhibit uncoordinated locomotion (Figure 5A). In fact, though these double mutants show more movement in the anterior half of their bodies than *nca-1(gf),* they propagate body waves to their posterior half even more poorly than the *nca-1(gf)* mutant. However, both *sek-1* and *nsy-1* partially suppress the loopy waveform of the *nca-1(gf)* mutant (Figure 5A, B) and in radial locomotion assays, *sek-1* and *nsy-1* weakly suppressed the *nca-1(gf)* locomotion defect (Figure 5C). Additionally, both *sek-1* and *nsy-1* partially suppress the small body size of *nca-1(gf)* (Figure S4A). Together these data suggest that mutations in the SEK-1 p38 MAPK pathway suppress some aspects of the *nca-1(gf)* mutant.

**Figure 5.**
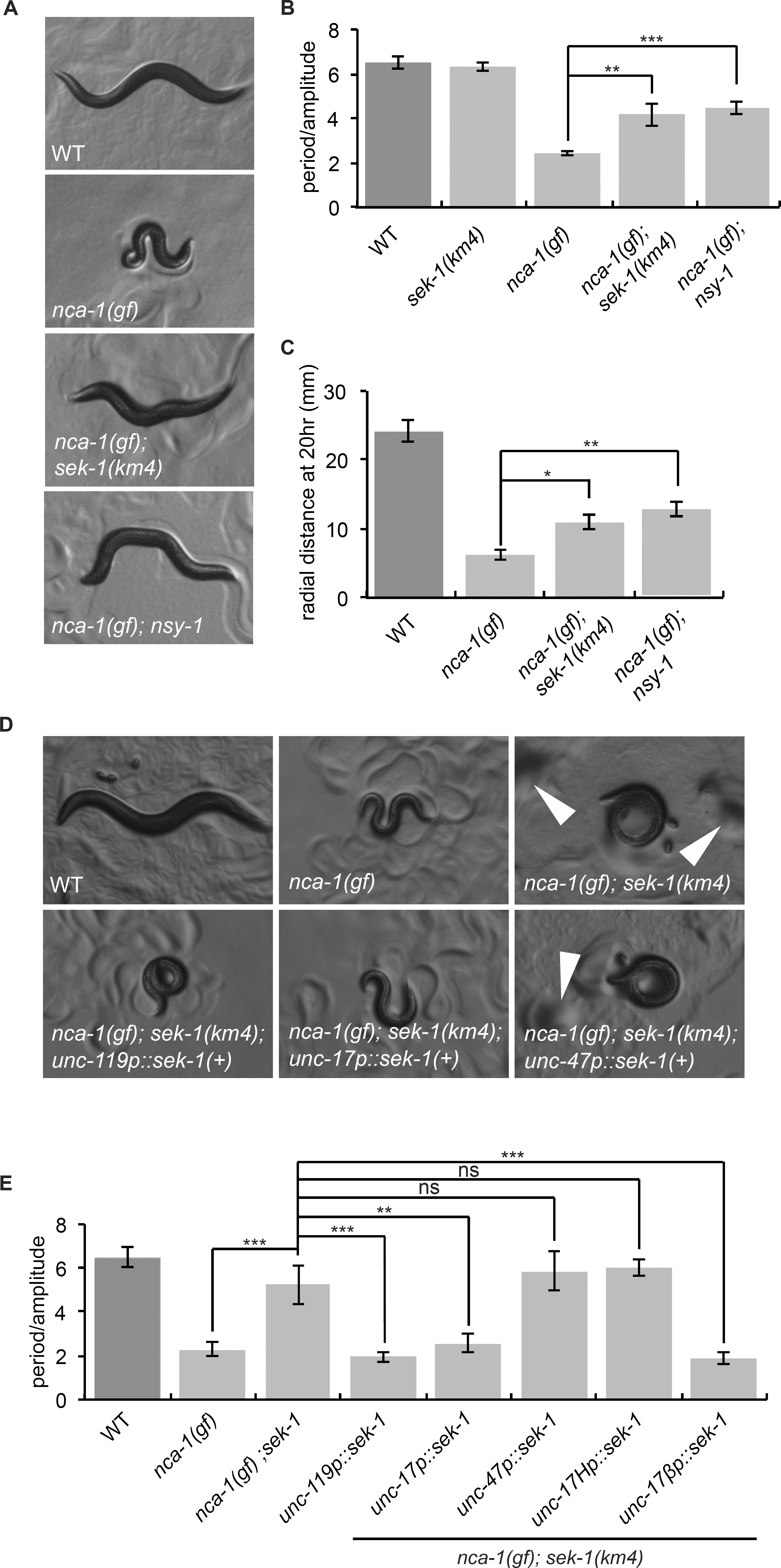
*sek-1* and *nsy-1* weakly suppress *nca-1(gf)* (A) *nca-1(gf)* mutants are small, loopy, and uncoordinated. The phenotypes of *nca-1(ox352)* animals are partially suppressed by *sek-1(km4)* and *nsy-1(ok593).* Photos of first-day adults. (B) *sek-1(km4)* and *nsy-1(ok593)* suppress the loopy waveform of *nca-1(gf).* ***, p<0.001; **, p<0.01. Error bars = SEM, n=5. (C) *sek-1* and *nsy-1* suppress *nca-1(gf)* locomotion. *nca-1(gf)* animals travel a small distance from the center of the plate in the radial locomotion assay. *nca-1(gf) nsy-1(ok593)* and *nca-1(gf) sek-1(km4)* worms move further than *nca-1(gf)* worms. **, p<0.01; *, p<0.05. Error bars = SEM, n=30. (D) *sek-1* acts in acetylcholine neurons to modulate the size and loopy waveform of *nca-1(gf). nca-1(ox352) sek-1(km4)* animals expressing *sek-1* in all neurons *(unc-119p::sek-1(+))* or in acetylcholine neurons *(unc-17p::sek-1(+))* are loopy and small like *nca-1(gf),* but *nca-1(ox352) sek-1(km4)* animals expressing *sek-1* in GABA neurons *(unc-47p::sek-1(+))* resemble *nca-1(gf) sek-1.* White arrowheads depict food piles created by *nca-1(gf) sek-1(km4)* animals due to their uncoordinated locomotion. Such food piles are not made by *nca-1(gf)* animals. (E) *sek-1* acts in acetylcholine motorneurons to modulate the loopy waveform of *nca-1(gf). nca-1(ox352) sek-1(km4)* worms expressing *sek-1* in all neurons *(unc-119p::sek-1(+)),* acetylcholine neurons *(unc-17p::sek-1(+)),* or acetylcholine motorneurons *(unc-17fip::sek-1(+))* are loopy like *nca-1(gf),* but *nca-1(ox352) sek-1(km4)* worms expressing *sek-1* in GABA neurons *(unc-47p::sek-1(+))* or head acetylcholine neurons *(unc-17Hp::sek-1(+))* are similar to *nca-1(gf) sek-1.* ***, p< 0.001, **, p<0.01; ns, p>0.05. Error bars = SEM, n=5.

Given that *sek-1* acts in acetylcholine neurons to modulate wild-type and *egl-30(tg26)* locomotion, we tested whether *sek-1* also acts in these neurons to suppress *nca-1(gf).* Expression of *sek-1* in all neurons or in acetylcholine neurons of *nca-1(gf) sek-1(km4)* animals restored the *nca-1(gf)* loopy phenotype (Figure 5D, E). By contrast, expression of *sek-1* in GABA neurons did not affect the loopy posture of the *nca-1(gf) sek-1* double mutant (Figure 5D, E). These data suggest that *sek-1* acts in acetylcholine neurons to modulate the body posture of *nca-1(gf)* as well. However, in radial locomotion assays, expression of *sek-1* in none of these neuron classes significantly altered the movement of the *nca-1(gf) sek-1* double mutant (Figure S4B), though the weak suppression of *nca-1(gf)* by *sek-1* in this assay makes it difficult to interpret these negative results. To further narrow down the site of action of *sek-1* for its NCA suppression phenotypes, we expressed it in subclasses of acetylcholine neurons. Surprisingly, expression of *sek-1* in acetylcholine motorneurons but not head acetylcholine neurons was sufficient to restore the loopy posture of the *nca-1(gf)* mutant (Figure 5E), the opposite of what we found for *sek-1* modulation of the loopy posture of the activated Gq mutant, suggesting that the loopy posture of *nca-1(gf)* mutants may result from excessive NCA-1 activity in acetylcholine motorneurons. Additionally, expression of *sek-1* in either the head acetylcholine neurons or the motorneurons restored the *nca-1(gf)* small body size phenotype (Figure S4C). We make the tentative conclusion that *sek-1* acts in acetylcholine neurons to modulate *nca-1(gf)* body posture and size, but we were not able to conclusively narrow down its site of action further, possibly due to the uncoordinated phenotype of *nca-1(gf)* and the weaker suppression of *nca-1(gf)* by *sek-1*.

## Discussion

The p38 MAPK pathway has been best characterized as a pathway activated by a variety of cellular stresses and inflammatory cytokines (Kyriakis and Avruch 2012), but it has also been implicated in neuronal function, including some forms of mammalian synaptic plasticity (Bolshakov *et al.* 2000; Rush *et al.* 2002; Huang *et al.* 2004). In this study we identified a new neuronal role for the mitogen-activated protein kinase kinase SEK-1 and the p38 MAPK pathway as a positive modulator of locomotion rate and Gq signaling. The physiological importance of this pathway is clear under conditions of elevated Gq signaling but is less obvious during normal wild-type locomotion, consistent with the observation that *sek-1* mutations have a relatively weak effect on synaptic transmission in a wild-type background (Vashlishan *et al.* 2008). Thus, the SEK-1 p38 MAPK pathway may be more important for modulation of Gq signaling and synaptic strength than for synaptic transmission per se.

In addition to SEK-1, we identified other p38 pathway components that modulate Gq signaling. Specifically, we found that *tir-1, nsy-1* and *pmk-1 pmk-2* mutants exhibit locomotion defects identical to *sek-1* and suppress activated Gq, suggesting that they act in a single p38 pathway to modulate signaling downstream of Gq. These results indicate a redundant function for PMK-1 and PMK-2 in modulating locomotion rate and Gq signaling. PMK-1 and PMK-2 also act redundantly for some other neuronal roles of the p38 pathway, such as the development of the asymmetric AWC neurons and to regulate induction of serotonin biosynthesis in the ADF neurons in response to pathogenic bacteria (Shivers *et al.* 2009; Pagano *et al.* 2015). By contrast, PMK-1 acts alone in the intestine to regulate innate immunity and in interneurons to regulate trafficking of the GLR-1 glutamate receptor (Pagano *et al.* 2015; Park and Rongo 2016).

What are the downstream effectors of the SEK-1 p38 MAPK pathway that modulates locomotion? There are several known downstream effectors of p38 MAPK signaling in *C. elegans*, including the transcription factor ATF-7 (Shivers et al. 2010). Our data indicate that ATF-7 is not required for the p38 MAPK-dependent modulation of Gq signaling. The p38 MAPK pathway may activate molecules other than transcription factors or may activate multiple downstream effectors.

How does the SEK-1 p38 pathway modulate the output of Gq signaling? One of the pathways that transduces signals from Gq includes the RhoGEF Trio/UNC-73, the small GTPase Rho, and the cation channel NALCN/NCA-1 (Williams *et al.* 2007; Topalidou *et al.* 2017). Compared to other pathways downstream of Gq, mutants in the Rho-Nca pathway are particularly strong suppressors of the loopy waveform phenotype of the activated Gq mutant (Topalidou *et al.* 2017). Similary, we found that mutations in the SEK-1 p38 MAPK pathway strongly suppress the loopy waveform of the activated Gq mutant, suggesting that the SEK-1 pathway might modulate Gq signal output through the Rho-Nca branch. Consistent with this, we found that mutations in the SEK-1 p38 MAPK pathway partially suppress an activated NCA-1 mutant. Given the precedence for direct phosphorylation of sodium channels by p38 to regulate channel properties (Wittmack *et al.* 2005; Hudmon *et al.* 2008), it is possible that PMK-1 and PMK-2 phosphorylate NCA-1 to regulate its expression, localization, or activity.

Consistent with the observation that Gq acts in acetylcholine neurons to stimulate synaptic transmission (Lackner *et al.* 1999), we found that *sek-1* acts in acetylcholine neurons to modulate the locomotion rate in both wild-type and activated Gq mutants. *sek-1* also acts in acetylcholine neurons to modulate the loopy waveform of both activated Gq and activated *nca-1* mutants, and the size of activated *nca-1* mutants. However, our data attempting to narrow down the site of action of *sek-1* suggest that it may act in both head acetylcholine neurons and acetylcholine motorneurons, and that the waveform is probably controlled by at least partially distinct neurons from those that control locomotion rate. Further work will be required to identify the specific neurons where Gq, NCA-1 and the SEK-1 pathway act to modulate locomotion rate and waveform, and determine whether they all act together in the same cell.

## Acknowledgements

We thank Dennis Kim and Chiou-Fen Chuang for strains, Pin-An Chen and Erik Jorgensen for the *nca-1(gf)* mutant *ox352,* Chris Johnson for the fine mapping of *yak42,* Jordan Hoyt for help with Galaxy to analyze WGS data, and Dana Miller for providing access to her microscope camera. Some strains were provided by the CGC, which is funded by NIH Office of Research Infrastructure Programs (P40 OD010440). J.M.H was supported in part by Public Health Service, National Research Service Award T32GM007270, from the National Institute of General Medical Sciences. M.A. is an Ellison Medical Foundation New Scholar. This work was supported by NIH grant R00 MH082109 to M.A.

## Supplementary Information

**Table S1.**
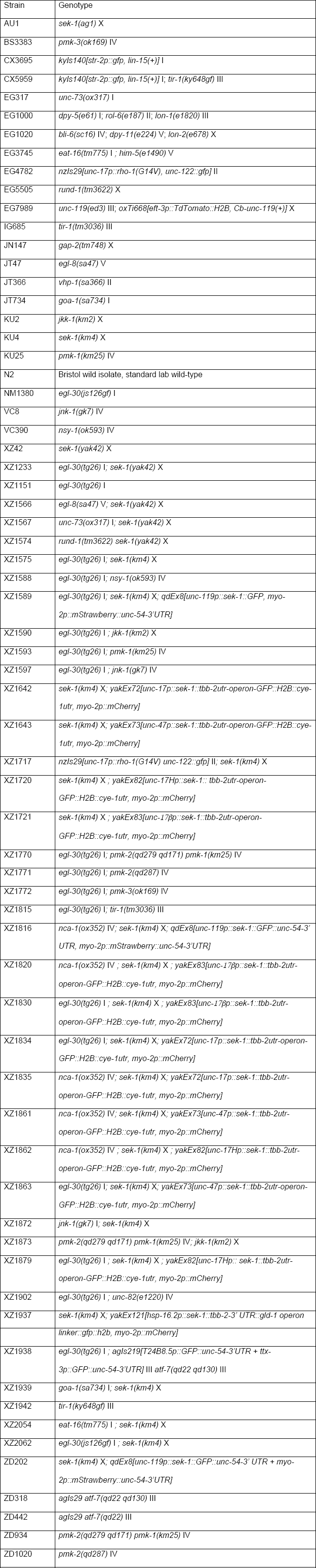
Strain List.

**Table S2.**
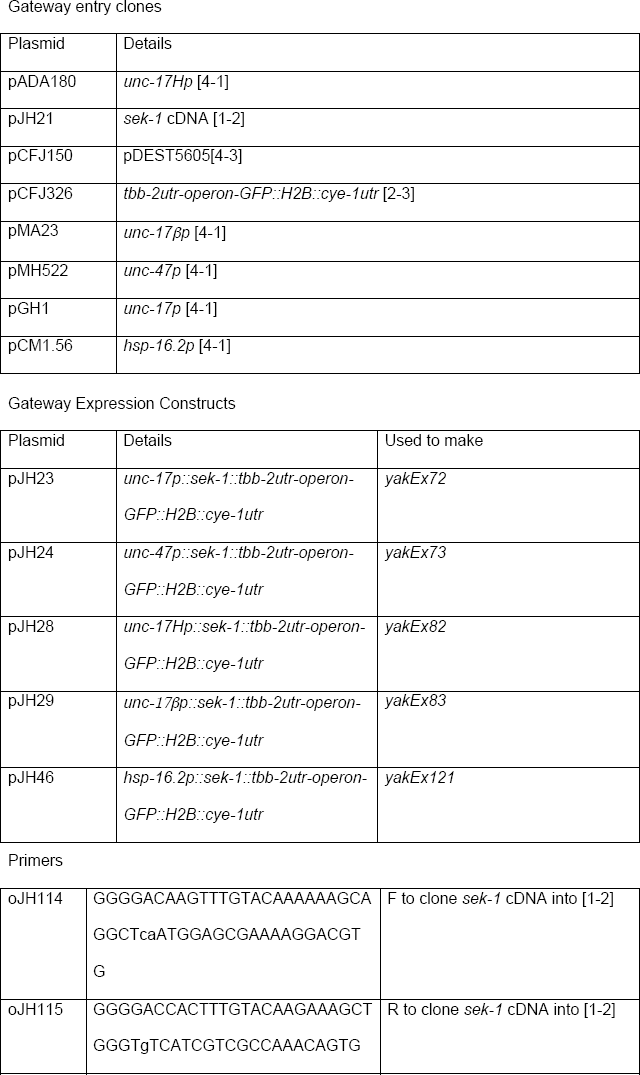
Plasmids and Primers.

**Table S3.** Statistical Tests.

| Figure | Test | p value |
| --- | --- | --- |
| 1C | One-way ANOVA and Bonferroni's Multiple Comparison Test<br>WT vs *sek-1(km4)* (ns) | < 0.001 |
|  | WT vs *egl-30(tg26)* (p<0.001)<br>*egl-30(tg26)* vs *egl-30(tg26); sek-1(km4)* (p<0.001)<br>*egl-30(tg26)* vs *egl-30(tg26); sek-1(yak42)* (p<0.001)<br>*egl-30(tg26)* vs *egl-30(tg26); unc-82(e1220)* (ns) |  |
| 1E | One-way ANOVA and Bonferroni's Multiple Comparison Test<br>WT vs *egl-30(tg26)* (p<0.001)<br>*egl-30(tg26)* vs *egl-30(tg26); sek-1(yak42)* (p<0.001)<br>*egl-30(tg26)* vs *egl-30(tg26); sek-1(km4)* (p<0.001) | < 0.001 |
| 1F | One-way ANOVA and Dunnett's Multiple Comparison Test<br>WT vs *sek-1(yak42)* (p<0.001)<br>WT vs *sek-1(km4)* (p<0.001) | < 0.001 |
| 1G | One-way ANOVA and Bonferroni's Multiple Comparison Test<br>WT vs *egl-30(js126)* (p<0.001)<br>WT vs *sek-1(km4)* (p<0.001)<br>*egl-30(js126)* vs *egl-30(js126); sek-1(km4)* (p<0.001) | < 0.001 |
| 1H | One-way ANOVA and Bonferroni's Multiple Comparison Test<br>WT vs *egl-30(js126)* (p<0.001)<br>*egl-30(js126)* vs *egl-30(js126); sek-1(km4)* (p<0.001) | <0.001 |
| 2A | One-way ANOVA and Bonferroni's Multiple Comparison Test<br>*sek-1(km4)* vs *sek-1(km4); qdEx8[unc-119::sek-1(+)]* (p<0.001) | < 0.001 |
| 2B | One-way ANOVA and Bonferroni's Multiple Comparison Test<br>*sek-1(km4)* vs *sek-1(km4); yakEx72[unc-17p::sek-1(+)]* (p<0.001)<br>*sek-1(km4)* vs *sek-1(km4); yakEx73[unc-47p::sek-1(+)]* (ns) | <0.001 |
| 2D | One-way ANOVA and Bonferroni's Multiple Comparison Test<br>WT vs *egl-30(tg26)* (p<0.001) | <0.001 |
|  | *egl-30(tg26); sek-1(km4)* vs<br>*egl-30(tg26); sek-1(km4); qdEx8[unc-119p::sek-1(+)]*<br>(p<0.001)<br>*egl-30(tg26); sek-1(km4)* vs<br>*egl-30(tg26); sek-1(km4); yakEx72[unc-17p::sek-1(+)]*<br>(p<0.001)<br>*egl-30(tg26); sek-1(km4)* vs<br>*egl-30(tg26); sek-1(km4); yakEx73[unc-47p::sek-1(+)]* (ns) |  |
| 2E | Kruskal-Wallis Test and Dunn's Multiple Comparison Test<br>*egl-30(tg26); sek-1(km4)* vs<br>*egl-30(tg26); sek-1(km4); qdEx8[unc-119p::sek-1(+)]* (p<0.001)<br>*egl-30(tg26); sek-1(km4)* vs<br>*egl-30(tg26); sek-1(km4); yakEx72[unc-17p::sek-1(+)]* (p<0.01)<br>*egl-30(tg26); sek-1(km4)* vs<br>*egl-30(tg26); sek-1(km4); yakEx73[unc-47p::sek-1(+)]* (ns) | < 0.001 |
| 2F | One-way ANOVA and Bonferroni's Multiple Comparison Test<br>*sek-1(km4)* vs *sek-1(km4); yakEx121[hsp-16.2p::sek-1(+)]*<br>(p<0.001) | < 0.001 |
| 3A | Unpaired t test, two-tailed | < 0.001 |
| 3B | One-way ANOVA and Bonferroni's Multiple Comparison Test<br>*egl-30(tg26)* vs *egl-30(tg26); tir-1(tm3036)* (p<0.001) | < 0.001 |
| 3C | Unpaired t test, two-tailed | < 0.001 |
| 3D | One-way ANOVA and Bonferroni's Multiple Comparison Test<br>*egl-30(tg26)* vs *egl-30(tg26); nsy-1(ok593)* (p<0.001) | < 0.001 |
| 3E | One-way ANOVA and Dunnett's Multiple Comparison Test | < 0.001 |
|  | WT vs *pmk-1(km25)* (ns)<br>WT vs *pmk-2(qd287)* (p<0.05)<br>WT vs *pmk-2(qd279 qd171) pmk-1 (km25)* (p<0.001)<br>WT vs *pmk-3(ok169)* (p<0.001) |  |
| 3F | One-way ANOVA and Dunnett's Multiple Comparison Test<br>*egl-30(tg26)* vs *egl-30(tg26); pmk-1(km25)* (p<0.001)<br>*egl-30(tg26)* vs *egl-30(tg26); pmk-2(qd287)* (p<0.001)<br>*egl-30(tg26)* vs *egl-30(tg26); pmk-2(qd279 qd171) pmk-1 (km25)* (p<0.001)<br>*egl-30(tg26)* vs *egl-30(tg26); pmk-3(ok169)* (p<0.001) | < 0.001 |
| 3H | One-way ANOVA and Bonferroni's Multiple Comparison Test<br>*egl-30(tg26)* vs *egl-30(tg26); tir-1(tm3036)* (p<0.001)<br>*egl-30(tg26)* vs *egl-30(tg26); nsy-1(ok593)* (p<0.001)<br>*egl-30(tg26)* vs *egl-30(tg26); sek-1(km4)* (p<0.001)<br>*egl-30(tg26)* vs *egl-30(tg26); pmk-2(qd279 qd171) pmk-1(km25)* (p<0.001) | < 0.001 |
| 4A | One-way ANOVA and Bonferroni's Multiple Comparison Test<br>*egl-8(sa47)* vs *egl-8(sa47); sek-1(yak42)* (p<0.001) | < 0.001 |
| 4B | One-way ANOVA and Bonferroni's Multiple Comparison Test<br>*sek-1(yak42)* vs *rund-1(tm3622); sek-1(yak42)* (p<0.001) | < 0.001 |
| 4C | One-way ANOVA and Bonferroni's Multiple Comparison Test<br>*unc-73(ox317)* vs *unc-73(ox317); sek-1(yak42)* (ns) | < 0.001 |
| 4E | One-way ANOVA and Bonferroni's Multiple Comparison Test<br>WT vs *nzIs29[unc-17p::rho-1(G14V)]* (p<0.001)<br>WT vs *nzIs29[unc-17p::rho-1(G14V)]; sek-1(km4)* (p<0.001) | < 0.001 |
|  | *nzls29[unc-17p::rho-1(G14V)]* vs<br>*nzls29[unc-17p::rho-1(G14V)]; sek-1(km4)* (ns) |  |
| 4F | One-way ANOVA and Bonferroni's Multiple Comparison Test<br>*nzls29[unc-17p::rho-1(G14V)]* vs<br>*nzls29[unc-17p::rho-1(G14V)]; sek-1(km4)* (p<0.001) | < 0.001 |
| 5B | One-way ANOVA and Bonferroni's Multiple Comparison Test<br>WT vs *sek-1(km4)* (ns)<br>WT vs *nca-1(ox352)* (p<0.001)<br>*nca-1(ox352)* vs *nca-1(ox352); sek-1(km4)* (p<0.01)<br>*nca-1(ox352)* vs *nca-1(ox352); nsy-1(ok593)* (p<0.001) | <0.001 |
| 5C | One-way ANOVA and Bonferroni's Multiple Comparison Test<br>WT vs *nca-1(ox352)* (p<0.001)<br>*nca-1(ox352)* vs *nca-1(ox352); nsy-1(ok593)* (p<0.01)<br>*nca-1(ox352)* vs *nca-1(ox352); sek-1(km4)* (p<0.05) | < 0.001 |
| 5E | One-way ANOVA and Bonferroni's Multiple Comparison Test<br>*nca-1(ox352)* vs *nca-1(ox352); sek-1(km4)* (p<0.001)<br>*nca-1(ox352); sek-1(km4)* vs<br>*nca-1(ox352); sek-1(km4); qdEx8[unc-119p::sek-1(+)]*<br>(p<0.001)<br>*nca-1(ox352); sek-1(km4)* vs<br>*nca-1(ox352); sek-1(km4); yakEx72[unc-17p::sek-1(+)]*<br>(p<0.01)<br>*nca-1(ox352); sek-1(km4)* vs<br>*nca-1(ox352); sek-1(km4); yakEx73[unc-47p::sek-1(+)]* (ns)<br>*nca-1(ox352); sek-1(km4)* vs | < 0.001 |
|  | *nca-1(ox352); sek-1(km4); yakEx82[unc-17Hp::sek-1(+)]* (ns)<br>*nca-1(ox352); sek-1(km4)* vs<br>*nca-1(ox352); sek-1(km4); yakEx83[unc-17βp::sek-1(+)]*<br>(p<0.001) |  |
| S1A | One-way ANOVA and Bonferroni's Multiple Comparison Test<br>WT vs *egl-30(tg26)* (p<0.01)<br>WT vs *sek-1(km4)* (p<0.001)<br>*egl-30(tg26)* vs *egl-30(tg26); sek-1(km4)* (p<0.001)<br>*sek-1(km4)* vs *egl-30(tg26); sek-1(km4)* (p<0.01) | < 0.001 |
| S1B | One-way ANOVA and Newman-Keuls Multiple Comparison Test<br>WT vs *sek-1(yak42)* (p<0.001)<br>WT vs *sek-1(ag1)* (p<0.001)<br>*sek-1(yak42)* vs *sek-1(ag1)* (ns) | < 0.001 |
| S1C | One-way ANOVA and Bonferroni's Multiple Comparison Test<br>WT vs *egl-30(tg26)* (ns)<br>*egl-30(tg26)* vs *egl-30(tg26); unc-82(e1220)* (p<0.001) | < 0.001 |
| S1D | One-way ANOVA and Bonferroni's Multiple Comparison Test<br>WT vs *sek-1(km4)* (p<0.001)<br>*goa-1(sa734)* vs *sek-1(km4)* (p<0.001)<br>*goa-1(sa734)* vs *goa-1(sa734); sek-1(km4)* (p<0.001) | < 0.001 |
| S1E | One-way ANOVA and Bonferroni's Multiple Comparison Test<br>WT vs *sek-1(km4)* (ns)<br>WT vs *goa-1(sa734)* (p<0.001)<br>*goa-1(sa734)* vs *goa-1(sa734); sek-1(km4)* (p<0.001) | < 0.001 |
| S1F | One-way ANOVA and Bonferroni's Multiple Comparison Test | < 0.001 |
|  | WT vs *sek-1(km4)* (p<0.001)<br>WT vs *eat-16(tm775)* (p<0.001)<br>*eat-16(tm775)* vs *eat-16(tm775); sek-1(km4)* (p<0.001)<br>*sek-1(km4)* vs *eat-16(tm775); sek-1(km4)* (ns) |  |
| S1G | One-way ANOVA and Bonferroni's Multiple Comparison Test<br>WT vs *eat-16(tm775)* (p<0.001)<br>*eat-16(tm775)* vs *eat-16(tm775); sek-1(km4)* (ns) | < 0.001 |
| S2A | One-way ANOVA and Bonferroni's Multiple Comparison Test<br>WT vs *sek-1(km4)* (p<0.001)<br>*sek-1(km4)* vs *sek-1(km4); yakEx82[unc-17Hp::sek-1(+)]* (p<0.05)<br>*sek-1(km4)* vs *sek-1(km4); yakEx83[unc-17βp::sek-1(+)]* (p<0.01) | < 0.001 |
| S2B | One-way ANOVA and Bonferroni's Multiple Comparison Test<br>*egl-30(tg26); sek-1(km4)* vs *egl-30(tg26); sek-1(km4); yakEx82[unc-17Hp::sek-1(+)]* (p<0.001)<br>*egl-30(tg26); sek-1(km4)* vs *egl-30(tg26); sek-1(km4); yakEx83[unc-17βp::sek-1(+)]* (p<0.001) | < 0.001 |
| S2C | One-way ANOVA and Bonferroni's Multiple Comparison Test<br>WT vs *egl-30(tg26)* (p<0.001 )<br>*egl-30(tg26); sek-1(km4)* vs *egl-30(tg26); sek-1(km4); qdEx8[unc-119p::sek-1(+)]* (p<0.001)<br>*egl-30(tg26); sek-1(km4)* vs *egl-30(tg26); sek-1(km4); yakEx72[unc-17p::sek-1(+)]* (p<0.001)<br>*egl-30(tg26); sek-1(km4)* vs | < 0.001 |
|  | *egl-30(tg26); sek-1(km4); yakEx73[unc-47p::sek-1(+)]* (ns)<br>*egl-30(tg26); sek-1(km4)* vs *egl-30(tg26); sek-1(km4); yakEx82[unc-17Hp::sek-1(+)]* (p<0.001)<br>*egl-30(tg26); sek-1(km4)* vs *egl-30(tg26); sek-1(km4); yakEx83[unc-17βp::sek-1(+)]* (ns) |  |
| S3A | One-way ANOVA and Bonferroni's Multiple Comparison Test<br>*jnk-1(gk7)* vs *jnk-1(gk7); sek-1(km4)* (p<0.01)<br>*jkk-1(km2)* vs *pmk-2(qd279 qd171) pmk-1 (km25); jkk-1(km2)* (p<0.05) | < 0.05 |
| S3B | One-way ANOVA and Bonferroni's Multiple Comparison Test<br>WT vs *egl-30(tg26)* (P<0.001)<br>*egl-30(tg26)* vs *egl-30(tg26); jkk-1* (ns)<br>*egl-30(tg26)* vs *egl-30(tg26); jnk-1* (ns)<br>*egl-30(tg26)* vs *egl-30(tg26); atf-7* (ns) |  |
| S3C | One-way ANOVA and Bonferroni's Multiple Comparison Test<br>WT vs *atf-7(qd22)* (p<0.001)<br>WT vs *atf-7(qd22 qd130)* (p<0.001) | < 0.001 |
| S3D | One-way ANOVA | p=0.806 |
| S3F | Unpaired t test, two-tailed | < 0.001 |
| S4A | One-way ANOVA and Bonferroni's Multiple Comparison Test<br>WT vs *nca-1(ox352)* (p<0.001)<br>*nca-1(ox352)* vs *nca-1(ox352); sek-1(km4)* (p<0.001)<br>*nca-1(ox352)* vs *nca-1(ox352); nsy-1(ok593)* (p<0.001) | <0.001 |
| S4B | Kruskal-Wallis Test and Dunn's Multiple Comparison Test | 0.001 |
|  | *nca-1(ox352);sek-1(km4)* vs *nca-1(ox352);sek-1(km4); qdEx8[unc-119p::sek-1(+)]* (ns)<br>*nca-1(ox352);sek-1(km4)* vs<br>*nca-1(ox352);sek-1(km4); yakEx72[unc-17p::sek-1(+)]* (ns)<br>*nca-1(ox352);sek-1(km4)* vs<br>*nca-1(ox352);sek-1(km4); yakEx73[unc-47p::sek-1(+)]* (ns) |  |
| S4C | One-way ANOVA and Bonferroni's Multiple Comparison Test<br>*nca-1(ox352)* vs *nca-1(ox352); sek-1(km4)* (p<0.001)<br>*nca-1(ox352); sek-1(km4)* vs<br>*nca-1(ox352); sek-1(km4); qdEx8[unc-119p::sek-1(+)]* (p<0.001)<br>*nca-1(ox352); sek-1(km4)* vs<br>*nca-1(ox352); sek-1(km4); yakEx72[unc-17p::sek-1(+)]* (p<0.001)<br>*nca-1(ox352); sek-1(km4)* vs<br>*nca-1(ox352); sek-1(km4); yakEx73[unc-47p::sek-1(+)]* (ns)<br>*nca-1(ox352); sek-1(km4)* vs<br>*nca-1(ox352); sek-1(km4); yakEx82[unc-17Hp::sek-1(+)]* (p<0.001)<br>*nca-1(ox352); sek-1(km4)* vs<br>*nca-1(ox352); sek-1(km4); yakEx83[unc-17βp::sek-1(+)]* (p<0.001) | <0.001 |

**Figure S1.**
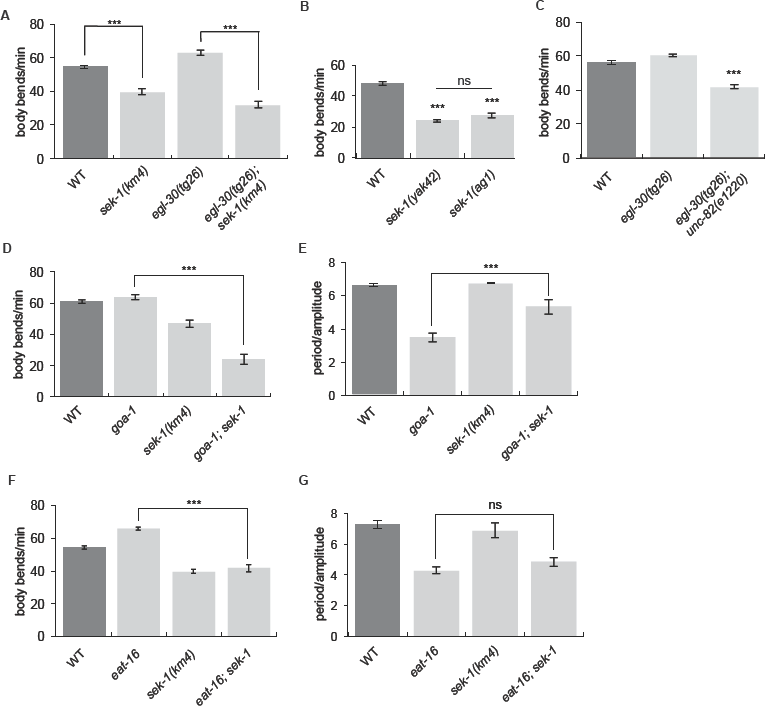
*sek-1* interacts with Gq and Go mutants. (A) The *sek-1(km4)* mutation suppresses the hyperactive locomotion of the activated Gq mutant *egl-30(tg26).* ***, p<0.001, error bars = SEM, n=20. (B) *sek-1(ag1)* mutant animals have slow locomotion. ***, p<0.001, error bars = SEM, n=10. (C) The *unc-82(e1220)* mutation reduces the locomotion rate of the activated Gq mutant *egl-30(tg26).* ***, p<0.001, error bars = SEM, n= 20. (D) *sek-1(km4)* suppresses the hyperactivity of *goa-1(sa734).* ***, p<0.001, error bars = SEM, n=20. (E) *sek-1(km4)* suppresses the loopy waveform of *goa-1(sa734).* ***, p<0.001, error bars = SEM, n=5. (F) *sek-1(km4)* suppresses the hyperactivity of *eat-16(tm775).* ***, p<0.001, error bars = SEM, n=20. (G) *sek-1(km4)* does not suppress the loopy waveform of *eat-16(tm775).* ns, p>0.05, error bars = SEM, n=5.

**Figure S2.**
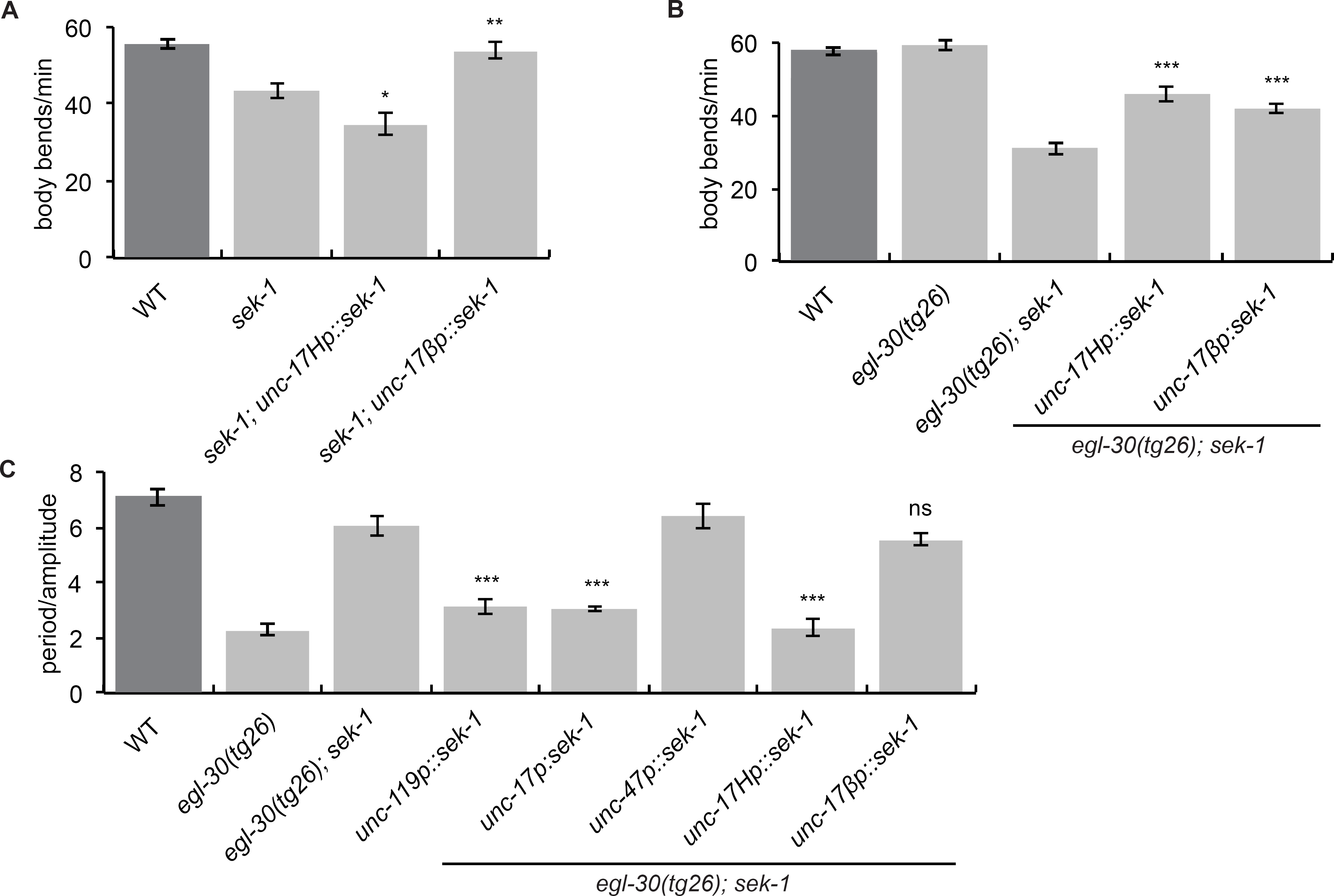
*sek-1* acts in both head acetylcholine neurons and acetylcholine motorneurons. (A) *sek-1* acts in acetylcholine motorneurons to modulate locomotion rate. The *sek-1* WT cDNA driven by the *unc-17fi* acetylcholine motorneuron promoter [*unc-17fip::sek-1(+)*] rescues the slow locomotion phenotype of *sek-1(km4)* worms, but *sek-1* expression in head acetylcholine neurons using the *unc-17H* promoter [*unc-17Hp::sek-1(+)*] does not rescue. **, p< 0.01; *, p<0.05 compared to *sek-1.* Error bars = SEM, n=20. (B) *sek-1* acts in both head acetylcholine neurons and acetylcholine motorneurons to modulate the locomotion rate of the activated Gq mutant *egl-30(tg26). egl-30(tg26) sek-1(km4)* worms expressing either *unc-17Hp::sek-1(+)* or *unc-17fip::sek-1(+)* have an increased locomotion rate compared to *egl-30(tg26) sek-1.* ***, p<0.001 compared to *egl-30(tg26) sek-1.* Error bars = SEM, n=20. (C) *sek-1* acts in head acetylcholine neurons to modulate the loopy waveform of the activated Gq mutant *egl-30(tg26). egl-30(tg26) sek-1(km4)* worms expressing *unc-17Hp::sek-1(+)* are loopy like *egl-30(tg26),* but *egl-30(tg26) sek-1(km4)* worms expressing *unc-17fip::sek-1(+)* are similar to *egl-30(tg26) sek-1.* ***, p<0.001; ns, p>0.05 compared to *egl-30(tg26) sek-1.* Error bars = SEM, n=5.

**Figure S3.**
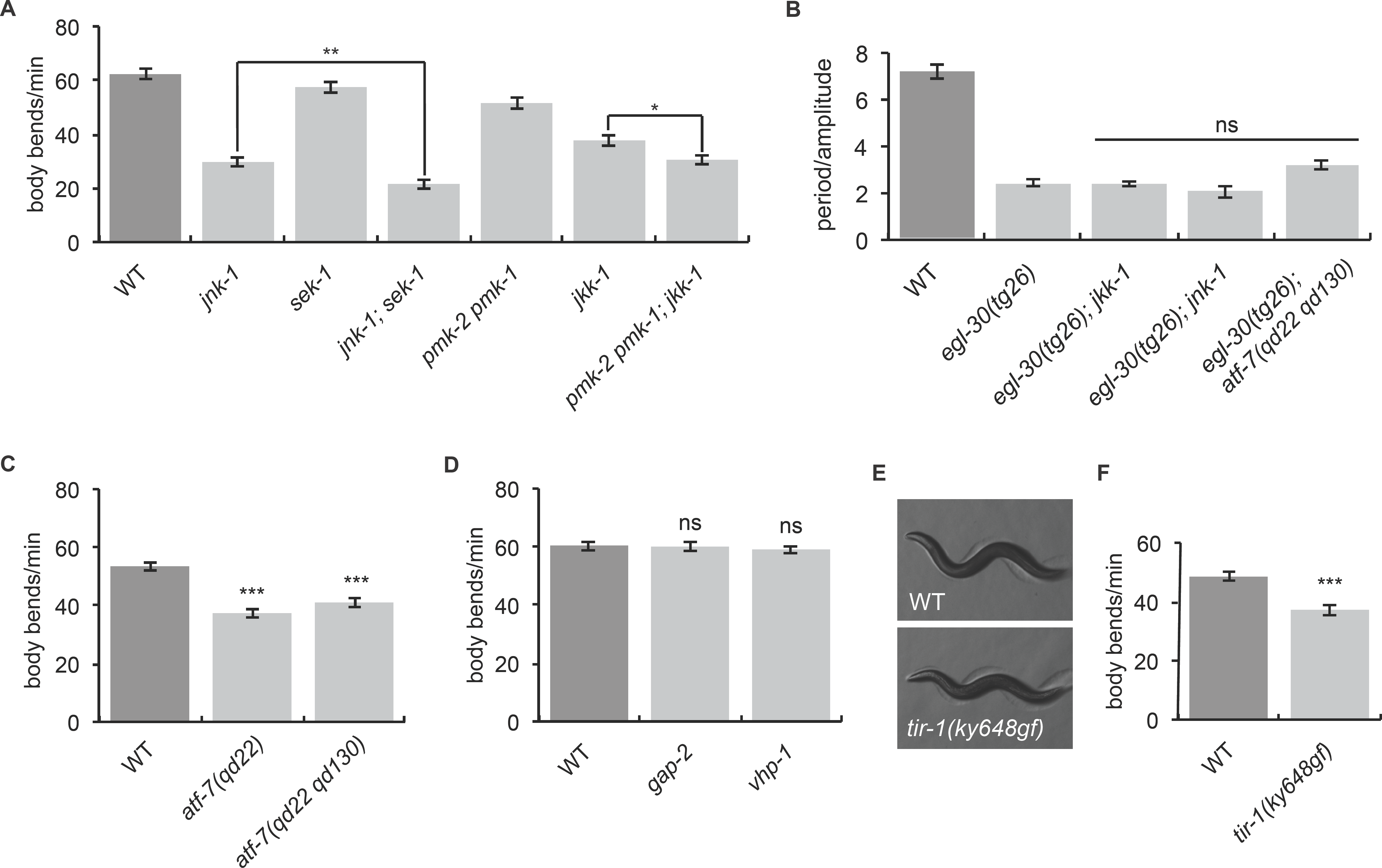
Locomotion of p38 and JNK MAPK pathway mutants. (A) *jkk-1* and *jnk-1* act in parallel to *sek-1* and *pmk-2 pmk-1.* The *jnk-1(gk7) sek-1(km4)* double mutant and *pmk-2(qd279 qd171) pmk-1(km25) jkk-1(km2)* triple mutants move more slowly than the respective individual mutants. **, p< 0.01, *, p<0.05. Error bars = SEM, n=20. (B) Mutations in *jkk-1, jnk-1,* and *atf-7* do not suppress the loopy waveform of the activated Gq mutant *egl-30(tg26).* ns, p>0.05, error bars = SEM, n=5. (C) Worms with gain-of-function or loss-of-function alleles of *atf-7* are slower than wild-type worms. ***, p< 0.001, error bars = SEM, n=20. (D) Worms lacking *gap-2* and *vhp-1* move like wild-type worms. Neither *gap-2(tm478)* nor *vhp-1(sa366)* confers a slow locomotion phenotype. ns, p>0.05 compared to WT. Error bars = SEM, n=20. (E-F) *tir-1(ky648gf)* animals do not have loopy or hyperactive locomotion. *tir-1(ky648gf)* worms have wild-type posture and are slower than wild-type animals. ***, p< 0.001, error bars = SEM, n=20.

**Figure S4.**
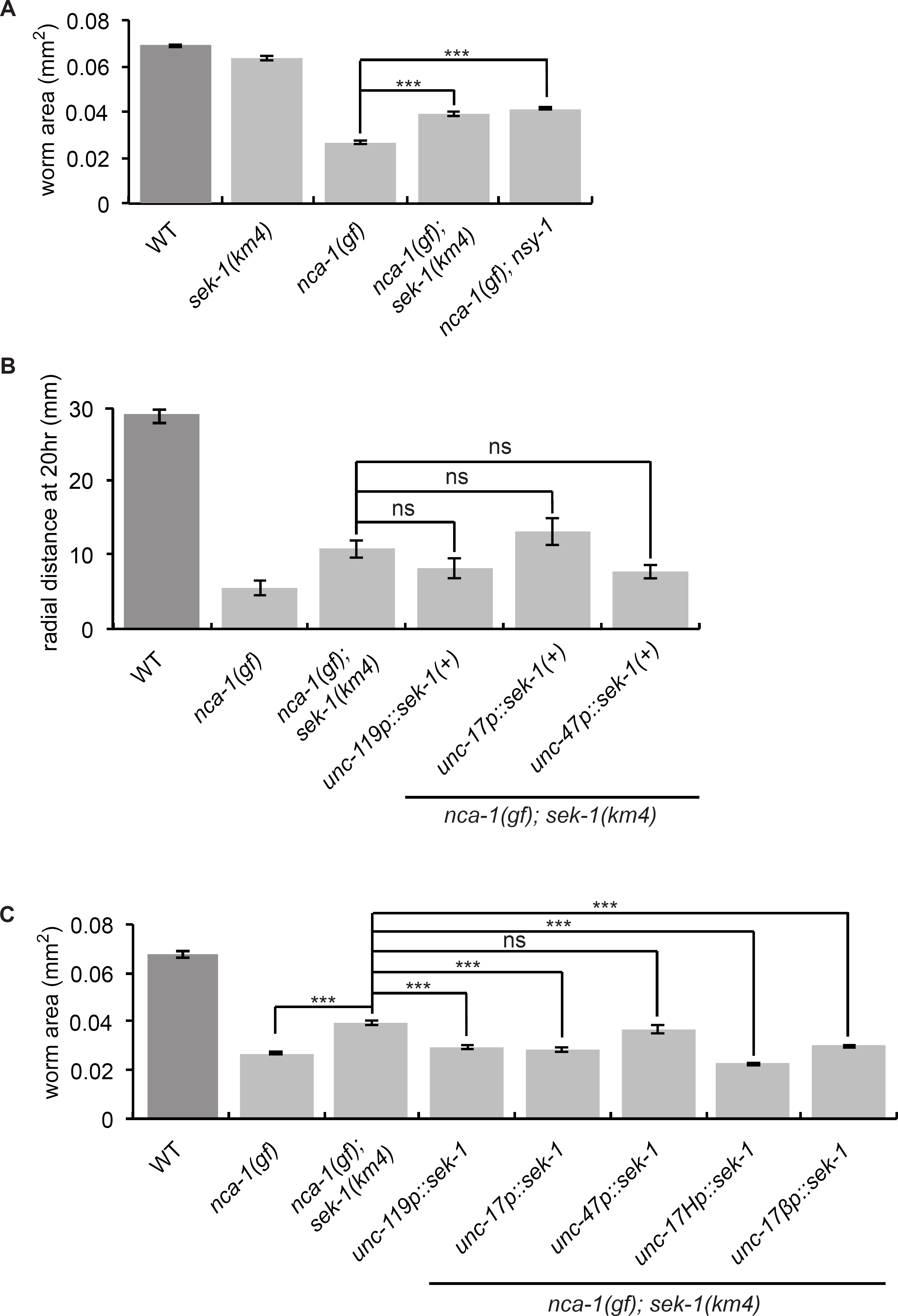
*sek-1* and *nsy-1* weakly suppress *nca-1(gf)* (A) Mutations in *sek-1* and *nsy-1* suppress the small body size of *nca-1(ox352)* mutant worms. ***, p<0.001, error bars = SEM, n=10. (B) None of the neuronal *sek-1* rescuing constructs reverse the radial locomotion phenotype of *nca-1(gf) sek-1(km4)* animals. ns, p>0.05. Error bars = SEM, n=19-24. (C) *sek-1* acts in both head acetylcholine neurons and acetylcholine motorneurons to control the body size of *nca-1(gf). nca-1(ox352) sek-1(km4)* worms expressing *sek-1* in all neurons *(unc-119p::sek-1(+)* acetylcholine neurons *(unc-17p::sek-1(+)),* head acetylcholine neurons *(unc-17Hp::sek-1(+)),* or acetylcholin motorneurons *(unc-17fip::sek-1(+))* have a similar size to *nca-1(gf),* but *nca-1(ox352) sek-1(km4)* worms expressing sek-1 in GABA neurons *(unc-47p::sek-1(+))* are similar to *nca-1(gf) sek-1.* ***, p< 0.001; ns, p>0.05. Error bars = SEM, n=7-10.

